# Machine learning-based detection of label-free cancer stem-like cell fate

**DOI:** 10.1101/2022.04.27.489645

**Authors:** Alexis J. Chambost, Nabila Berabez, Olivier Cochet-Escartin, François Ducray, Mathieu Gabut, Caroline Isaac, Sylvie Martel, Ahmed Idbaih, David Rousseau, David Meyronet, Sylvain Monnier

## Abstract

Most imaging methods rely on labelling biological samples in order to provide specific and easily detectable features. However, label-free imaging is a non-invasive and non-toxic alternative that requires accurate image analysis algorithm based on cell morphology. Such analysis has to deal with a high image variability while fewer features are extractable, so far, fast analysis of label-free brightfield microscopy (LFBM) images remains a challenging task. With the development of microfabricated devices during the last decades, high throughput image generation makes it possible to use machine learning-based algorithms in order to analyse LFBM images. Fast algorithms are also crucially needed to analyze high throughput experiments. In this paper, we provide a data-driven study in order to assess the complexity of LFBM time-lapses monitoring isolated cancer stem-like cells (CSCs) fate in non-adherent conditions. We combined for the first time individual cell fate and cell state temporality analysis in a unique algorithm. Several image analysis algorithms of increasing processing capacities were tested: a classical computer vision algorithm (CCVA), a shallow learning-based algorithm (SLBA) and a deep learning-based algorithm (DLBA). We show that our optimized DLBA has by far the best accuracy compared to CCVA and SLBA, is at least as accurate as other state-of-the-art DLBAs while being faster. With this study, we demonstrate that optimizing our DLBA accordingly to the image analysis problem can overall provide better results than pretrained models. Such a fast and accurate DLBA is therefore compatible with the generation of high throughput data and opens the route for on-the-fly analysis of CSC fate from LFBM time-lapses.

## Introduction

Labelling of living biological samples is a very powerful tool for studying molecules of interest, enhance contrast or increase microscopic resolution and has contributed to many groundbreaking advances in biology over the past decades^1^. However, labelling can also cause biochemical-induced artefacts such as protein overexpression which might alter cell metabolism^2^, phototoxicity which can affect cell behaviors^3^ and reconstruction processes from confocal images which can create artefacts^4^. For the specific identification of cell types, including multipotent cells such as cancer stem-like cells (CSCs) considered in this article, a combination of multiple markers^5^ is often required and therefore cannot be easily implemented in living cells and even less for patient-derived samples. Given these limitations cells morphological characteristics evaluated using label-free brightfield microscopy (LFBM) remains an excellent alternative, but the lack of specificity and contrast requires the development of dedicated image analysis tools.

In order to cope with these issues, several studies have used image analysis algorithms to provide quantified data from LBFM acquired images. For example, many classical computer vision algorithms (CCVA), *i*.*e*. based on a few sets of expert selected features, have been developed so far, such as thresholding methods for cell and vesicle segmentation^6^, or intensity projection from z-stacks for macrophages segmentation^7^. However, in the context of LFBM, there is a high sample-to-sample variance. Moreover, classical handcrafted methods based on few features and fixed decision rules require advanced programming skills and long development^8,9^.

High throughput data generation from microfabricated devices has made it possible to use machine learning-based algorithms for image analysis^10^ and notably analysis of LFBM acquired images. With shallow learning based methods, the decision rules can be data driven instead of handcrafted by experts. Shallow learning-based algorithms (SLBA) have been developped for several purposes, *e*.*g*. support vector machine has been trained in order to segment bacteria from z-stack images^11^ or random forest classifier has been used to quantify neurite outgrowth^12^. With deep learning-based algorithms (DLBA), the features can even be learned jointly with the decision rules in an end-to-end data driven fashion^13^. DLBAs are revolutionary since they considerably reduce the time of algorithm development. The power of DLBAs in deciphering the complexity of image analysis problems has already been proved on similar topics. Convolutional neural networks (CNNs) have been trained on LFBM acquired images with the aim of controlling focal position during time-lapses^14^, for counting the number of cells in microfluidic generated droplets^15^ and for quantifiying spheroid formation from single cells expressing a fluorescent reporter protein^16^. DLBAs have been shown to be more efficient than CCVAs for white blood cell classification^17^, and in a classification problem of multi-cellular spheroids, CNNs have outperformed SLBA when trained on a large data set^18^. Along the same line, improving feature extraction thanks to transfer learning from pre-trained CNNs can also enhance histopathological biopsy classification^19^.

In this article, we propose a full DLBA pipeline for the high-throughput prediction of cancer cells status imaged with LFBM i.e. division, death or quiescence. CSCs have the ability to escape cell death (anoïkis) while isolated and cultured in non-adherent conditions^20^. Such non-adherent conditions promote stemness and are classically used to maintain CSC populations.In this peculiar culture condition, CSCs are caracterized by their hability to divide and aggregate in multiple cell structures called tumorspheres similar to neural stem cell neurospheres.^21^ The proportion of CSCs within a given cell population derived from either tumor samples or *in vitro* cell cultures can therefore be evaluated by functional tests that are based on their capacity to survive as single cell and/or reconstitute a population of cancer cells *in vitro*^22^. Therefore, in order to provide time-resolved characterization of CSCs populations at individual cell level, we have generated data sets from LFBM time-lapses of patient-derived brain CSCs in suspension, cultured in micro-wells of a microfabricated chip recapitulating in this way classical stem cell culture conditions (Figure 1)^23,24^. This system can generate high-throughput images of micro-wells (*circa* 15 micro-wells per second) and requires fast and accurate analysis of micro-wells content to assess cell fate.

**Figure 1.**
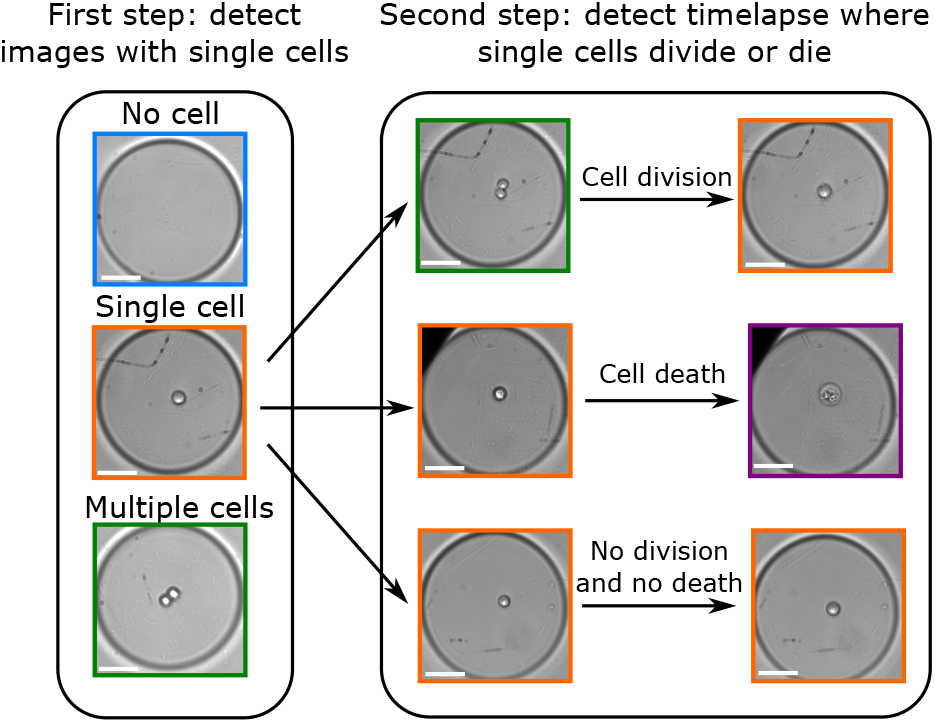
Visual abstract of the image analysis problem. First step: detection of single cells from 2D LFBM acquired images. Second step: when a single cell was detected, the corresponding LFBM 96-hour time-lapse was analysed in order to detect cell division or cell death. Examples of cell division with images of cytokinesis before daughter cells closely aggregate together, and of dead cells are provided. Scale bar: 50µm.

Here, we provide a data driven study to automatically process LFBM images of suspended cells. Despite a trivia appearance, the detection of these cell fates is challenging as i) cells can exhibit a large variety of shapes, ii) cells can be out-of-focus since they are cultured in non-adherent conditions, iii) these images contain less features than the ones available in mostly used image data sets like ImageNet^25^. For this purpose, we have developed several image analysis algorithms that can detect single cells and dynamically track their fate.

## Results

### Deep learning-based algorithm

We consider a problem of cell fate prediction. In informational terms, this is a classification invariant to translation. We therefore developed a convolutional neural network (CNN) adapted to a classification problem. We trained our CNN on data set which contains 17378 annotated images of glioblastoma-derived N14-0510 CSCs imaged with a 20X magnification every 30 minutes, a temporal resolution selected according to the proliferation characteristics of N14-0510 cell lineage in this culture condition. This model was based on successive layers of convolution and max pooling before a first fully connected layer together with a second fully connected layer that classified images as “Singles”, “Multiples”, “Death” or “Empty” (Figure 2a and Supplemental Figures S1a and S2a). Classification based on brightfield images was controled using fluorescent markers for cell numbers and cell death (Supplemental Figures S1b-c).

**Figure 2.**
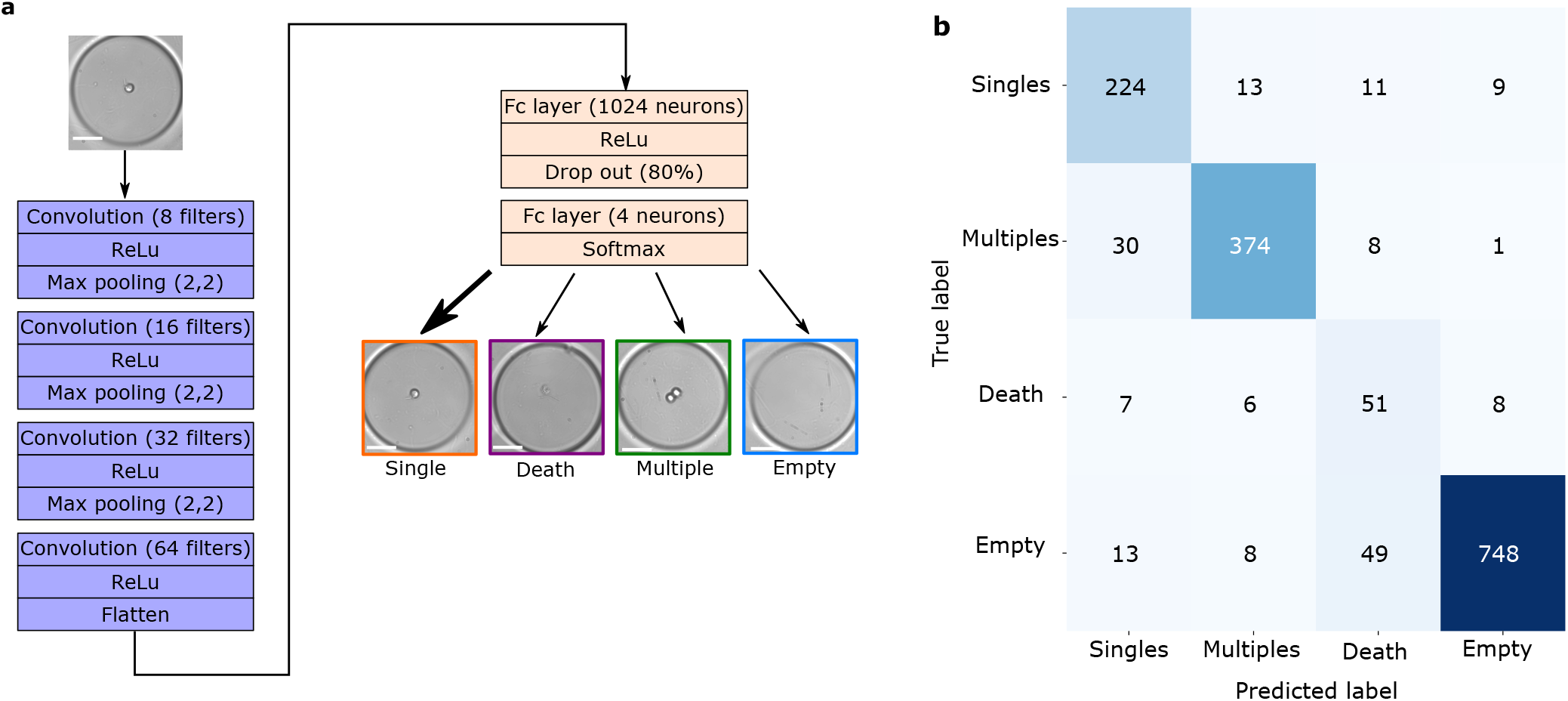
Optimised convoluted neural network (CNN). (a) Model summary of optimised CNN: 4 convolution layers before classification of images between “Singles”, “Multiples”, “Death” and “Empty”. Scale bars show 50µm. (b) Double entry table analysing differences between the ground truth (“True label”, vertical axis) and the prediction (“Predicted label”, horizontal axis). Optimized CNN provides accuracy of 91.2 *±* 0.17%. Recall and precision were respectively 87% and 81% for “Singles”, 90% and 93% for “Multiple”, 70% and 42% for “Death”, and 91% and 97% for “Empty”. Color dark stands for amount in images.

First, we have optimized the number of layers, the number of filters per convolution layer and the kernel size: four convolution layers of 8, 16, 32 and 64 filters respectively, and a kernel size of 15×15 pixels gave the best accuracy (Supplemental Figure S2c). Secondly, we have determined the optimal number of neurons in the first fully connected layer: 1024. Because of the high number of neurons in the first fully connected layer, a 80% drop out rate provided the best accuracy (Supplemental figures S2d-e). Finally, computing *circa* 8 million features (Supplemental Table S1), our CNN reached an accuracy of 91.2% (±0.17%) (Figure 2b). More precisely, precisions for “Singles”, “Multiples” and “Empty” classes reached 81.7%, 93.2% and 97.6% respectively. Nevertheless, some images were misclassified, notably for “Singles” and “Death” classes (Supplemental Figures S2f).

Our optimized CNN has also been compared to other cutting edge CNNs (Table 1). With the same purpose, Anagnostidis et al^15^ developed a simple similar CNN which provided a lower accuracy when applied on test data set: 82% (±0.26%). We have also trained pre-trained CNN on our training data set. Notably, Eulenberg et al^26^ developed an InceptionV3 based CNN in order to classify cell cycle phases. Although transfer learning-based algorithms were computing more features and required more computation time, they did not show better accuracy than our optimized CNN when applied on test data set (although VGG16 gives close results 91.2% (±0.13%)).

**Table 1.** Comparison of CCVA, SLBA, our optimized CNN and other publicly available CNNs. For the CNNs, we show the mean accuracy ± standard deviation after three model trainings.The mean of computation time per image is given in seconds ± standard deviation.

| Algorithm | Machine learning | Deep learning | Transfer learning | Number of features | Accuracy | Computation time per image (s) |
| --- | --- | --- | --- | --- | --- | --- |
| CCVA | — | — | — | 5 | 65% | $0.15 \pm 0.1$ |
| SLBA | + | — | — | 8 | 72% | $281 \pm 36$ |
| Classical CNN<sup>15</sup> | + | + | — | 162,900 | $82 \pm 0.26\%$ | $0.02 \pm 0.03$ |
| ResNet50<sup>27,28</sup> | + | + | + | 23,587,712 | $61.3 \pm 1.53\%$ | $0.06 \pm 0.07$ |
| InceptionV3<sup>26</sup> | + | + | + | 21,810,980 | $78.6 \pm 0.06\%$ | $0.09 \pm 0.13$ |
| VGG16<sup>29</sup> | + | + | + | 17,943,108 | $91.2 \pm 0.13\%$ | $0.05 \pm 0.04$ |
| Our CNN | + | + | — | 8,541,700 | $91.2 \pm 0.17\%$ | $0.02 \pm 0.03$ |

### Classical computer vision algorithm

To offer a baseline comparison with our optimized CNN, we also investigated our classification problem with a classical computer vision algorithm (CCVA). We developped a CCVA that pre-processed images in a few steps (Figure 3a-b): i) objects were segmented according to intensity threshold, ii) the largest and smallest segmented objects were filtered depending on their size in pixels, iii) the remaining objects were fitted to an ellipse, iv) according to the area and the roundness of the ellipse, the objects were classified as “Singles”, “Multiples”, “Death” or “Empty” if no object was segmented. Five features were analysed by CCVA: i) Intensity thresholding was based on Otsu’s method^30^, ii) size filtering has been empirically optimized based on 175 images of validation data set (Figures S2a-b), iii) position, as bordering objects were removed, iv) roundness and v) area thresholds were optimized among a wide range of values because they provided the best accuracy (Supplemental Figure S3a). When applied to test data set (Supplemental Figures S2a-b and Supplemental Table S2), CCVA only provided an accuracy of 65% (Figure 3c). More precisely, it appeared that a lot of images with single or multiple cells were classified as “Empty” (*circa* 57%), either because they were not segmented by intensity thresholding, or their size did not match the size filter. We figured out that there was a high image-to-image variability based on cell morphology, micro-well shape and depth of field (Supplemental Figure S3b). This variability explains why our CCVA with so few features (5) provided a low accuracy.

**Figure 3.**
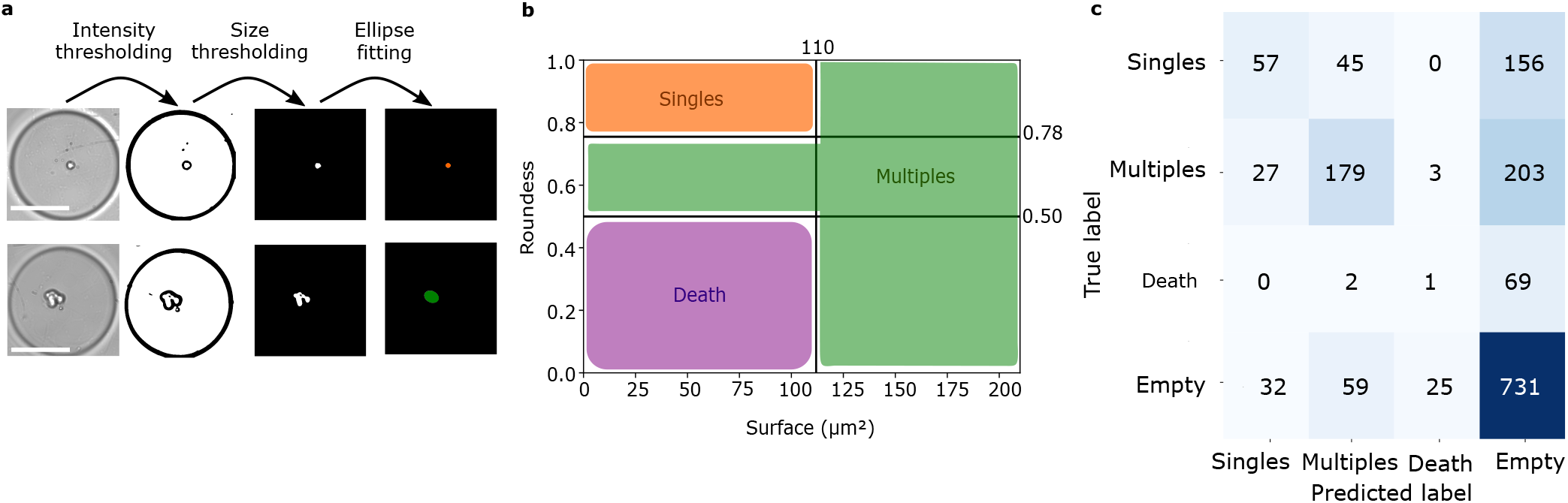
Classical computer vision algorithm (CCVA). (a) Images were processed through an intensity threshold before a size filtering. Remaining objects were fit to an ellipse. The first row was an image of a single cell and the second row depicted an image with multiple cells. Scale bars show 100µm. (b) According to area and roundness of the ellipse, images were classified as “Singles”, “Multiples”, “Death” or “Empty”. (c) Double entry table analysing differences between the ground truth (“True label”, vertical axis) and the prediction (“Predicted label”, horizontal axis). CCVA provides an accuracy of 65%. Recall and precision were respectively 22% and 66% for “Singles”, 42% and 62% for “Multiple”, 1% and 3% for “Death”, and 89% and 63% for “Empty”. Color dark stands for amount in images.

### Shallow learning-based algorithm

As machine learning-based algorithms can tackle image-to-image variability by training on large data sets, we wonder if shallow learning-based algorithm (SLBA) can cope with the complexity of our problem. Random forest has been shown to be one of the most accurate SLBA^31^. Here, we used a random forest classifier implemented under ilastik software^32^. Thanks to its pixel classification combined with object classification methods, several studies used this software for image analysis^12^. Based on these studies, we performed a pixel classification step by training the software to segment cells from the background on 40 images from validation data set (Supplemental Figures S2a-b and Supplemental Table S2). Then, using an object classification method, segmented objects were classified between “Singles”, “Multiples”, “Death” or “Empty” (Figure 4a). SLBA was processing respectively 5 and 3 features for pixel classification and object classification steps (Supplemental Table S3). Applied to the test data set (Supplemental Table S2), SLBA performed a better classification than CCVA: its accuracy reached 72% (Figure 4b). While many images with cells were still classified as “Empty” (*circa* 13%), there were also a lot of misclassifications between “Singles” and “Multiples” classes (*circa* 16%).

**Figure 4.**
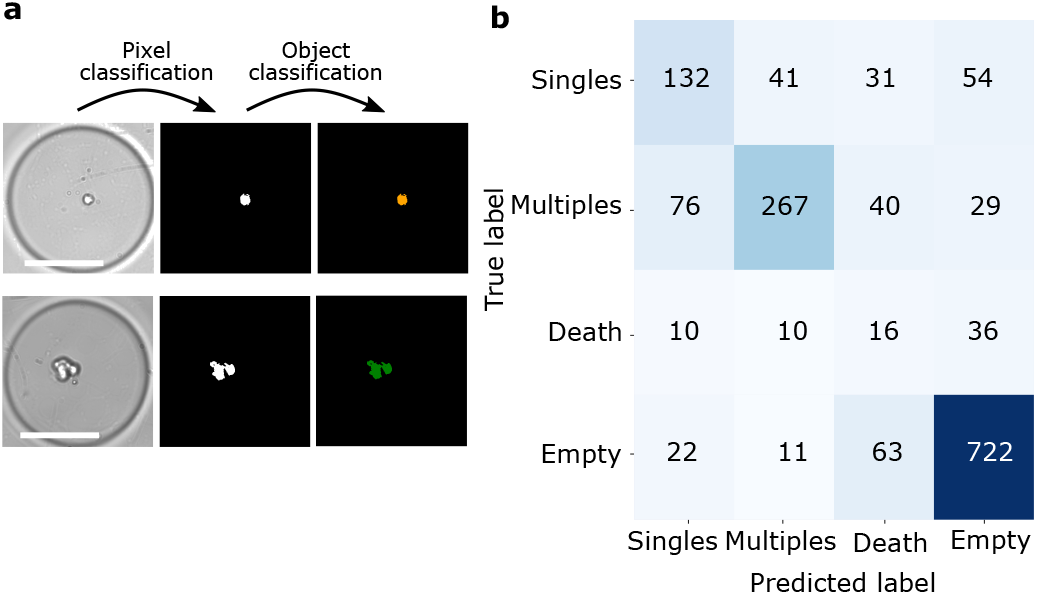
Shallow learning-based algorithm (SLBA). (a) The first step of pixel classification segments cells from the background. Then, thanks to an object classification tool, images were classified between “Singles”, “Multiples”, “Death” and “Empty” categories. The first row shows a single cell that was classified as a single cell and the second row depicted an image of multiple cells. Colour code between classes was shown in Fig. 2b. Scale bars show 100µm. (b) Double entry table analysing differences between the ground truth (“True label”, vertical axis) and the prediction (“Predicted label”, horizontal axis). SLBA provides an accuracy of 72%. Recall and precision were respectively 51% and 55% for “Singles”, 64% and 81% for “Multiple”, 22% and 10% for “Death”, and 88% and 85% for “Empty”. Color dark stands for amount in images.

Hence, we showed here that classification of images containing single cells, multiple cells, dead cells or no cell is a tough issue. It required to process a lot of parameters trained on large data sets, not only to find cells in the image but also to count them. Our optimized CNN was a more accurate and a much faster (computation time *circa* 60 ms per image) classifier than our CCVA and SLBA (Table 1 and Supplemental Figure S4). Nevertheless, our CNN’s misclassification led us to take into account the temporal dimension of our cell fate detection commitment.

### Temporal information encoding

As our algorithm analysed only 2D images, temporal information of time-lapse acquisition was lost. For example, two cells can adhere tightly to each other in suspension, while having the morpholgy of a single cell, but earlier events such as cytokinesis can demonstrate that two cells were actually present in the micro-well earlier. Considering the temporal encoding could therefore rescue our CNN’s misclassifications. In order to cope with this issue, we have developed a post-analytical decision tree (Figure 5). The detection of single cells, cell divisions and cell death with this decision tree was evaluated by computation of recall and precision on fully annotated LFBM time-lapses. It appeared that this method was very efficient to detect single cells (recall: 93%; precision: 96%), cell divisions (recall: 67%; precision: 94%) and cell death (recall: 82%; precision: 90%) from Time-lapse data set 1 (Table 2 and Supplemental Table S4). The impact of the frame-rate on the detection of divisions of our DLBA has also been investigated with Time-lapse data set 1: decreasing frame rate induced lower recall but precision was not modified (Supplemental Table S5). The probability thresholds and the majority voting can be considered as hyper-parameters that have been empirically tuned. Besides, we have not seen any single cell escaping from its micro-well during time-lapses annotation. Hence, we have not considered classification of image as “Empty” during the time-lapse of a micro-well if a single cell was detected at the beginning of the acquisition.

**Table 2.** Recall and precision for the detection of time-lapses with one single cell, of cell divisions and of cell death. The table shows results in percentage and absolute numbers for N14-0510 cells at 20X and 10X magnifications, and for N14-1525 cells at 20X and 10X magnifications (respectively 1091, 1179, 717 and 596 time-lapses analysed).

|  | Magnification | N14-0510 |  | N14-1525 |  |
| --- | --- | --- | --- | --- | --- |
|  |  | Recall | Precision | Recall | Precision |
| Single cell | 20X | 93% (281/301) | 96% (281/292) | 80% (142/176) | 92% (142/153) |
|  | 10X | 93% (312/334) | 99% (312/314) | 86% (138/160) | 97% (138/142) |
| Division | 20X | 67% (55/81) | 94% (55/58) | 79% (23/28) | 88% (23/26) |
|  | 10X | 80% (57/71) | 93% (57/61) | 93% (29/31) | 87% (29/33) |
| Cell death | 20X | 82% (96/117) | 90% (96/106) | 54% (27/50) | 79% (27/34) |
|  | 10X | 70% (60/85) | 89% (60/67) | 65% (36/55) | 92% (36/39) |

**Figure 5.**
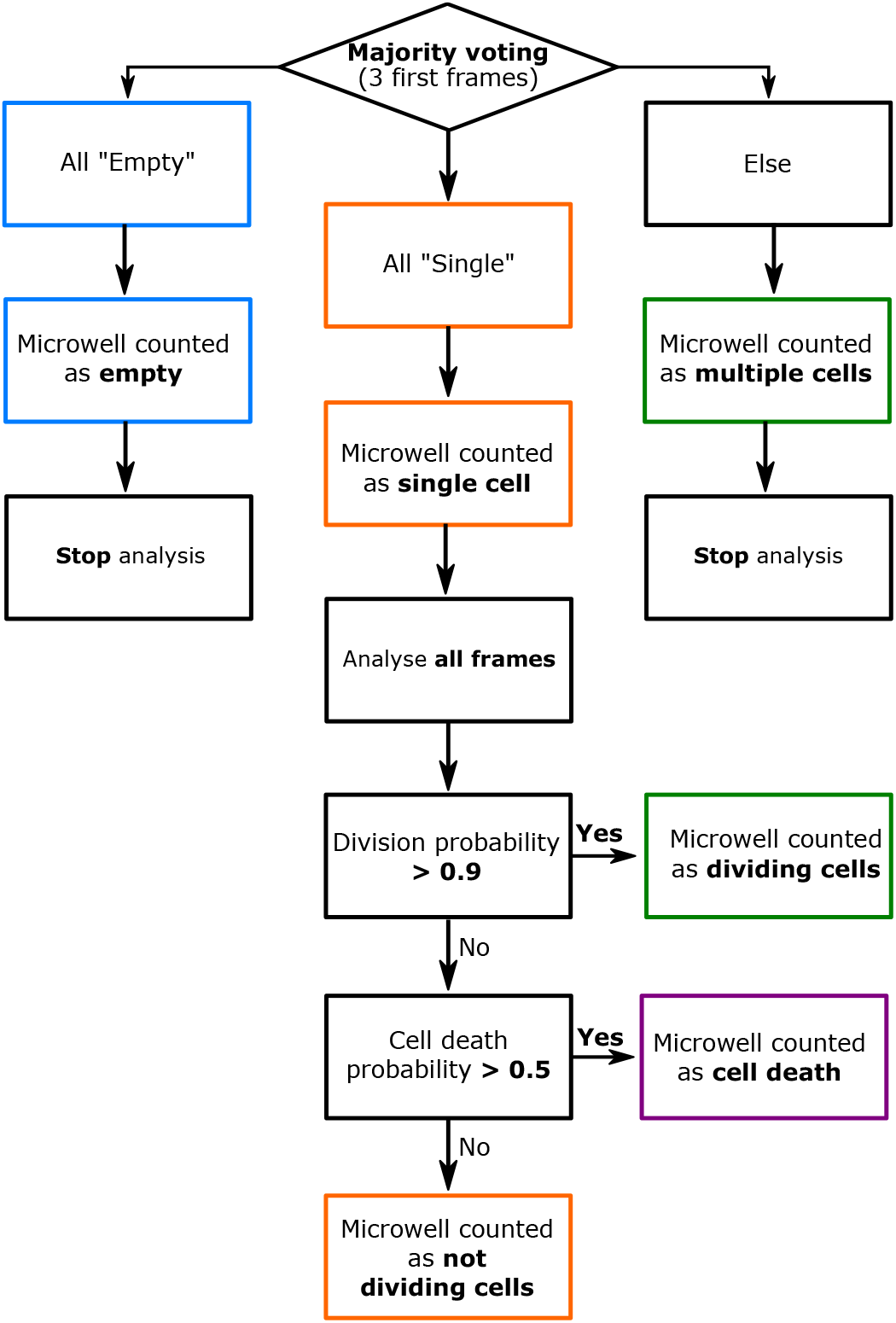
Downstream of the CNN, a decision tree considers the temporal information of cells time-lapses in order to detect single cells, cell divisions and cell death. First, we performed a majority voting on the first three frames of a micro-well. If all these three frames were classified as “Empty”, the micro-well was considered as containing no cell and its time-lapse was not further analysed. Then, if any of these three first frames was classified as “Multiple” or “Death”, the micro-well was considered as containing multiple cells or dead cells and was discarded from the analysis too. Finally, if all the three first frames were classified as “Single”, the micro-well was considered as containing a single cell and the analysis kept going. Among each time-lapse with an initial single cell, if two frames were classified as “Multiple” with a >0.9 probability (of which one >0.99), we considered that a cell division occurred in the micro-well. Else, if any frame was classified as “Death” with a >0.5 probability, we considered that a cell death occurred in the micro-well. In any other case, we considered that there were still a living but non-dividing cell in the micro-well.

### Impact of imaging magnification

As our annotated data set only contains images of N14-0510 cells acquired with a 20X magnification, we investigated the robustness of our algorithm to detect single cells, cell divisions and cell death of N14-0510 cells imaged at a 10X magnification. In order to assess the effect of a lower spatial resolution (Supplemental Figure S5), we annotated the Time-lapse data set 2 (Supplemental Table S4). Although detecting cell death was less sensitive with the 10X magnification compared to the 20X one (69% *versus* 82%), it remains as precise (89% (10X) *versus* 90% (20X)). Recall (93% (10X) *versus* 93% (20X)) and accuracy for single cell detection were similar (99% (10X) *versus* 96% (20X)) (Table 2). Surprisingly, cell divisions were even more sensitive at the 10X magnification (80%) compared to 20X (67%) while the precision remained similar (93% (10X) *versus* 93% (20X)).

### Testing our DLBA on another CSC line

Although our DLBA was trained on N14-0510 CSCs, we challenged it on the Time-lapse data sets 3 and 4, which were respectively composed of images of N14-1525 cells with 20X and 10X magnifications (Supplemental Figure S5 and Supplemental Table S4). N14-1525 cells were morphologically more heterogeneous than N14-0510 cells as *circa* 23% of N14-1525 had a much larger size (supplemental Figure S5 and Suppelemental Table S6). Results are summed up in the Table 2. Recalls for the N14-1525 cells were overall slightly decreased compared to N14-0510 cells, but precisions were similar. So, we considered that the global performance remained excellent and we showed that our DLBA could be used on different cellular models.

### Dynamics of cell divisions and cell death using our DLBA

As we compared the accuracy of our DLBA to other CNNs and assessed its robustness to different magnifications and brain CSC models, we next tested its ability to extract biologically relevant data such as time-resolved division and death rates. We performed 4 experiments with N14-0510 cells, with 3 replicates each. Time-lapses were LFBM acquired for 96 hours, with a 20X magnification objective, using an 80-minute imaging interval. After 4 days, time-lapses were analysed with our DLBA. The analysis of each experiment took less than 1 hour. In total, 2780 single cells were detected and 17% (±5%) underwent cell division within 96 hours (Figure 6a). When looking at the dynamic curve of cell divisions, we saw that most divisions occurred before 48 hours, with a curve flattening after 60 hours. Because the low number of late divisions (after 60 hours) was overwhelmed by initial number of single cells, computing dynamics did not allow a clear view of the latest divisions. In order to emancipate from the initial number of single cells, we have plotted the instantaneous division rate (Figure 6b). The initial division rate was *circa* 0.03 division/hour, but interestingly, it was decreasing over time. Regarding cell death, 38% (±9%) of single cells died during the 96-hour time-lapse (Figure 6c). Dynamics of cell death seemed rather linear, suggesting that instantaneous death rate should be relatively constant. Indeed, instant death rate was constant *circa* 0.03 death/hour, but it seemed to increase after 72 hours (up to 0.07 death/hour) (Figure 6d). Taken together, these results show that our DLBA can track automatically single cell fate and follow dynamics of cell phenotype over time at high throughput rate.

**Figure 6.**
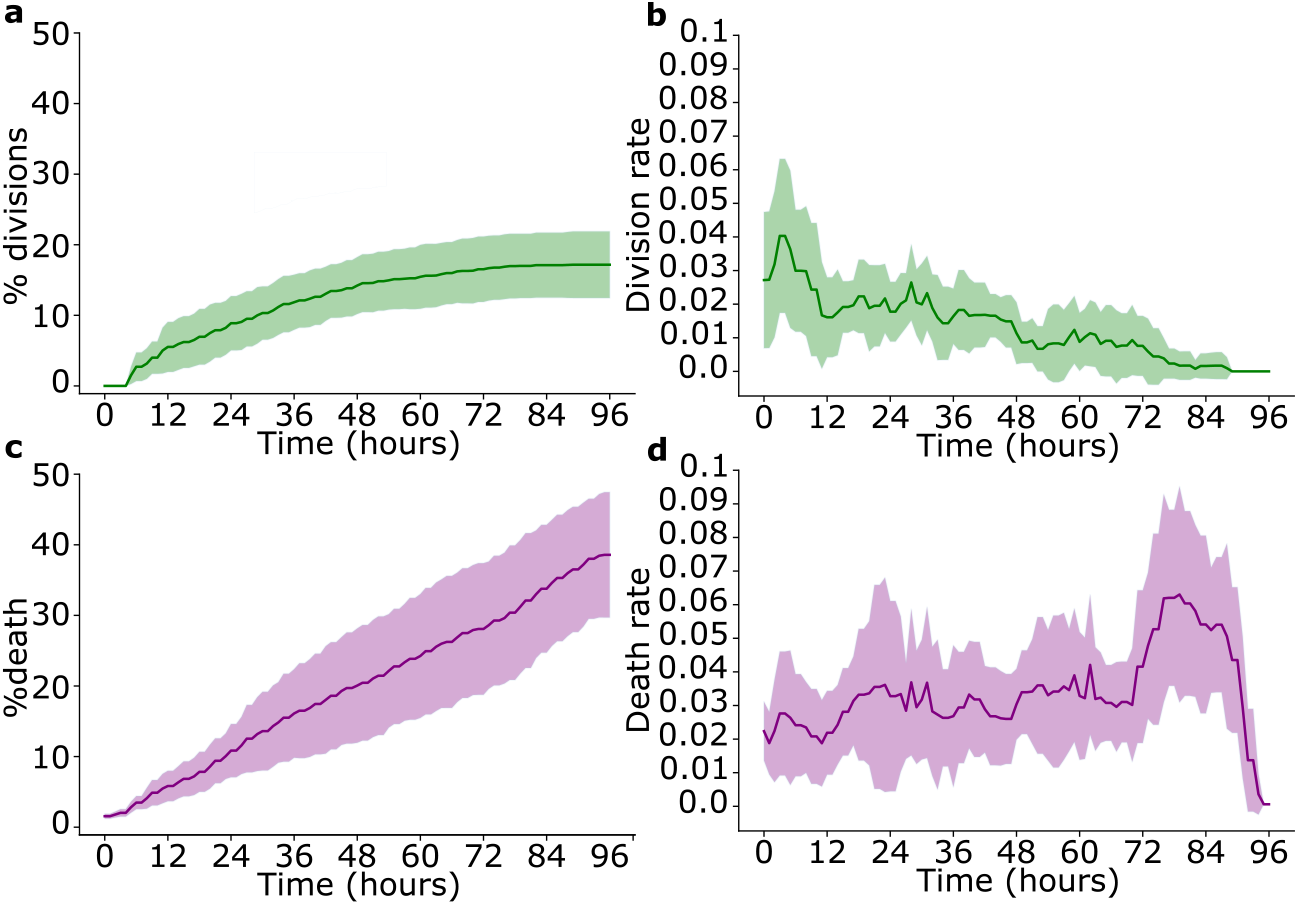
N14-0510 cell phenotyping analysis by our DLBA during LFBM time-lapse. Cumulative dynamics of cell divisions (a) and cell death (c) were computed over time. Relative division rate (b) and death rate (d) were computed, over sliding 6-hour temporal windows. Plots show means and standard deviations, over 4 experiments with three replicates each (2780 time-lapses analysed in total).

## Discussion

Biological samples can exhibit an important sample-to-sample variability, which may either reflect an heterogeneity of morphology or originate from the imaging methods themselves^9^. The apparent lack of specificity and the fluctuations of contrast can turn the analysis of LFBM images into a complex challenge. In addition, cells maintained in non-adherent culture conditions can frequently get out of focus, by convection or cell mobility, which affects their contrast and therefore our abilities to accurately detect cells. We propose here for the first time an efficient and precise method to evaluate simultaneously individual cell fate of CSCs (division, death, quiescence) including temporal information encoding of cell dynamics. Moreover, we show here that segmentation-based approaches are limited for our cell classification problem and that optimizing a classical CNN according to the problem can provide results as good as pretrained large models and save computational time/power.

Indeed, CCVA could not cope with the experimental variability and thus provided low accuracy (65%). While giving slightly better results in terms of cell segmentation, SLBA’s accuracy remains fairly low (72%). This might be due to a low number of features used to distinguish between the “Single” and “Multiples” classes. While the processing time for the SLBA could be largely optimized, with the parameters used, it takes approximately 5 minutes per image of 600×600 pixels and consequently can hardly be applied to high throughput generated data. We show here that CNNs seem to be the most accurate and faster algorithms for such image classification problem. Over 8 million features were processed by our optimized CNN that reached an accuracy of 91.2%. Indeed, only up to 2% of the error can be attributed to human misclassification (Supplemental Figure S1b-c), which confirms that the classification problem described here is indeed very challenging. In addition, our optimized CNN and the VGG16 pre-trained CNN provided similar results, suggesting that we almost reach the intrinsic complexity of our classification problem.

The complexity of our image data sets can be compared to other similar already published image analysis studies. With the correlation of DNA content and label-free morphological features, Blasi et al developed a SLBA in order to classify cell images at different steps during the cell cycle, but the precision for mitosis detection remained low (*circa* 45%)^33^. More recently, with the same purpose, this team combined LFBM images and fluorescent stains and extracted more features from this new data set, so their DLBA provided a better q precision for mitosis detection (*circa* 70%)^26^. With the approach of cell counting Anagnostidis et al developed a CNN that was very accurate for counting polyacrylamide beads but its accuracy was a bit lower when counting cells (*circa* 85%)^15^. Indeed, cell images exhibited more complex morphological features than polyacrylamide bead images. Concerning the detection of dead cells, Riba et al developed a DLBA in order to differentiate viable cells from dead cells. Interestingly, similarly to our results, they showed that their optimised CNN was more accurate (*circa* 80%) than more complex networks^34^. Here, our DLBA cannot only detect images of micro-wells with single cells with high recall and precision (93% and 96% respectively), but it can also detect dividing and dead cells with high reliability (i.e. recall and precision at 20X magnification are respectively 67% and 94%; and 82% and 90%).

Pre-trained CNNs have already provided excellent results to detect cell death or differentiation for adherent cells on a 2D surface^27,29,35^. Interestingly, such CNNs did not provide a better accuracy than ours (91.2%) while requiring larger computing times (See Table 1). Transfer learning CNNs have been pre-trained on the publicly available ImageNet database^25^. While it has already been reported that transfer learning did not always improve deep learning performances^36^, other results suggested that CNNs pre-trained on images dedicated to a similar purpose enhance network accuracy^19^. Statistics of natural images are known to produce in the Fourier domain power law spectrum in log-log scales^37^. This means there are no specific size of object, but rather objects of all sizes. In our images instead, we have a single object with a given size, typically a blob which will produce a spectrum with holes (zeros) in the Fourier spectrum. This may explain the relative inefficiency of the pre-trained models on our problem. Therefore, by comparison with this most related literature, the imaging and the image analysis have been optimized in order to deliver biologically relevant results with large statistics and high throughput. Indeed each micro-well contains a low number of cells or no cells at the beginning of the experiment. Consequently, the complexity of the informational task is reduced and this accounts for the much simpler and smaller neural network that we obtained as best solution. Therefore, the use of the micro-well systems not only increases the experimental throughput but also reduces the computational complexity of the neural network. From a methodological point of view, our CNN architecture is demonstrated to provide better results than standard architectures with a low number of hyperparameters to be tuned. Our approach enables easy time dependent processing and is more compatible with on-the-fly analysis. This is obtained thanks to the joint optimization of hardware (micro-well approach) and software (small CNN) which results in a globally more efficient solution.

Although our DLBA provided good predictions for N14-1525 cell images whereas it has been trained on N14-0510 images, we should consider adding images of other cell lines in the training database. Indeed, we found out that single N14-1525 cells which had a larger diameter compared to the average cell diameter observed in training images of N14-0510 cells were misclassified as “Multiple” (Supplemental Figure S5 and Supplemental Table S6). An efficient way to improve performances of the CNN could be adding to our image data sets more images from different cell lines with various shapes or textures, for example by adding cell models derived from other patients or tissues. Although this annotating step require some more work, it would be still less efforts than programming a new CCVA optimised to each new cell line. Another very interesting perspective would be implementing temporal convolution to our CNN in order to better take into account the temporal dimension of time-lapses. Despite implying the re-annotation of the training datasets used so far, the analysis of time series by CNNs should allow to significantly improve the prediction of cell fate^38^, for example by predicting cell divisions before it happens based on cell morphological characteristics.

We have also investigated the dynamics of cell division and death of CSCs originated from glioblastoma. These glioblastoma CSCs are at the centre of controversies because we are still missing reliable molecular marker to specifically identify them^22^. Morever, recent single cell transcriptomic data suggest that glioblastoma tumor-forming cells are rather defined by a continuum of differentiation than by clear distinctions between CSCs and differentiated cancer cells populations^39^–41. Patel et al suggested that CSCs express transcripts of cell cycle related genes at low level, and therefore that these cells likely have low division rates. This heterogeneity between slow and fast dividing cells could be recapitulated in Fig. 6b and might cells that are unequally positioned on the continuum of differentiation phenotypes. Besides, CSCs have been shown to escape anoïkis^42^, and more specifically, N14-0510 cells have been reported to display an increased expression of anti-apoptotic factors under non-adherent culture conditions^43^. Inasmuch our microfabricated chip prevents cell-substrate adhesion, cells that were still alive at the end of the time-lapse might have stem cell properties. Finally, coupling our DLBA to a micro-fabricated device dedicated to drug screening would provide a relevant image analysis pipeline in order to assess, on-the-fly and at high throughput rate, drug effects on cell divisions, cell death and CSCs fate modifications.

## Methods

### Microfabricated device

Manufacturing process has already been detailed in Goodarzi et al^23,24^. Briefly, 200µm diameter micro-wells were molded with 2% agarose solution over PDMS counter-moulds. Agarose was then immobilized on (3-Aminopropyl)triethoxysilane coated glass cover-slips. Eventually, GSCs were seeded in the agarose micro-wells.

### Cell lines and culture

N14-0510 and N14-1525 cell lines were kindly provided by A.I. and M.G. labs and were derived from diagnosed WHO grade IV glioblastoma before established as cellular models maintain in non-adherent conditions. They were maintained under normoxia at 37°C in incubator, in Dulbecco’s modified Eagle’s medium/nutrient mixture F12 (Life, 31330-095) complemented with N2 (Life, 17502-048) at 1X, B27 (Life, 17504-001) at 1X, 100 U/ml penicillin-streptomycin (Life, 15140-122) and FGF2 (Miltenyi Biotec, 130-104-922), EGF (Miltenyi Biotec, 130-093-825) (20ng/mL both) and heparin 0.00002% (Sigma, H3149). Unless otherwise specified, cells were cultured in Ultra low attachment T75 flasks (ThermoFisher, Ultra Low Adherent, 10491623). Cells were passaged weekly with Accumax (Sigma, A7089) at a density of 600 000 cells in 20 ml of complete media. The medium was renewed twice a week and mycoplasma tests were regularly performed. Hoechst staining (Sigma, H6024) was used to control cell number (10-100 ng/mL) and TO-PRO-1 iodide (ThermoFicher, T3602) was used to control cell viability (1:50). Cell size measurements were performed fter enzymatic dissociation of tumorsphere as previously detailed, the cell suspensions were quantified using the automatic LUNA FL cell counter (Logos biosystems) following the manufacturer’s instructions. Briefly, cells were stained with acridine orange and propidium iodide stain solution and were immediatly imaged with both brightfield and dual fluorescence optics to discriminate dead from living cells, and to estimate the size of living cells.

### Image setup

Samples were images with a Leica DMIRB microscope. Microscope was located in a impervious box with 37°C controlled temperature (LIS Cube) and 5% CO2 air (LIS Brick gaz mixer). The camera (Andor Neo 5.5 SCMOS), shutter (Vincent associated D1) and stage (Prior proscan II) were controlled with micromanager 1.4.22^44^. The light source was provided by a LED (Thorlabs MWWHL4). 20X and 10X numerical aperture objectives were both from Leica.

### Hardwares and softwares

Computing was performed with Windows 10, 64 bit operating system, Intel(R) Core(TM) i7-7700 3.60GHz processor. GPU used was NVIDIA quadro p600. All scripts were written in Python 3.7^45^. Libraries used were Mahotas 1.4.11^46^, OpenCV 4.2.0^47^, Seaborn^48^, numpy^49^, SciPy^50^, pandas^51^, matplotlib^52^ and Tensorflow 2.3.0^53^. Version of Ilastik used was 1.3.2post1^32^. Pretrained CNN were found at https://www.tensorflow.org/api_docs/python/tf/keras/applications.

### Data sets and code availability

Annotated data set was composed of 17378 LFBM acquired images. Images have been manually annotated. All cells from this data set were N14-0510 cells imaged with a 20X magnification. Amount of images per class was: 2871 “Singles” images, 4615 “Multiples” images, 803 “Death” images and 9089 “Empty” images (Supplemental Figure S5a). The number and viability of cells seen in brightfield has been controlled by fluorescent microscopy (Supplemental Figure S1b-c). 10% of these images were randomly selected in order to generate a validation data set, and another 10% was also randomly selected for the test data set. Remaining images constitute the training data set, on which we performed data augmentation in order to balance number of images between the four classes (Supplemental Figure S2a-b). Parameters of SLBA and CCVA were optimized respectively with 40 and 175 images manually selected from validation data set (Supplemental Table S2). Performance of DLBA, SLBA and CCVA were all compared on test data set. Computation times were compared on 1, 10, 100 and 1000 images randomly selected from test data set. Time-lapses were performed with a time interval of 40 minutes (Supplemental Table S4), except for the dynamics of cell division and death (Figure 6) where a 30 minutes interval was used. Time-lapse data set 1 was composed of 1091 annotated time-lapses of N14-0510 cells imaged with 20X magnification. There were 356 empty micro-wells, 434 micro-wells with 2 cells or more, 301 micro-wells with single cells which 81 divided and 117 died. Time-lapse data set 2 was composed of 1179 annotated time-lapses of N14-0510 cells imaged with 10X magnification. There were 363 empty micro-wells, 482 micro-wells with 2 cells or more, 334 micro-wells with single cells which 71 divided and 85 died. Time-lapse data set 3 was composed of 717 annotated time-lapses of N14-1525 cells imaged with 20X magnification. There were 231 empty micro-wells, 310 micro-wells with 2 cells or more, 176 micro-wells with single cells which 26 divided and 34 died. Time-lapse data set 4 was composed of 596 annotated time-lapses of N14-1525 cells imaged with 10X magnification. There were 222 empty micro-wells, 214 micro-wells with 2 cells or more, 160 micro-wells with single cells which 31 divided and 55 died. Image databases and codes can be found at https://github.com/chalbiophysics/XXX.

### Statistics

Classical efficiency scores were performed to evaluate and compare algorithms. Those scores involve true positives (positive images correctly classified, or TP), true negatives (negative images correctly classified, or TN), false positives (negative images misclassified, or FP) and false negatives (positive images misclassified, or FN). Accuracy was computed when CCVA, SLBA and the various CNNs:

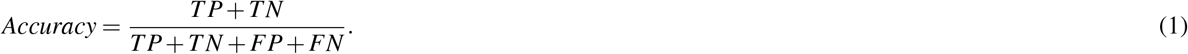

When assessing time-lapse classification and comparison between cell lines and magnifications by DLBA, recall and precision were computed:

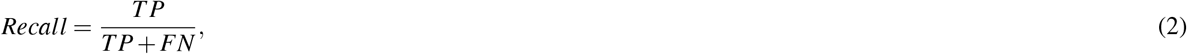

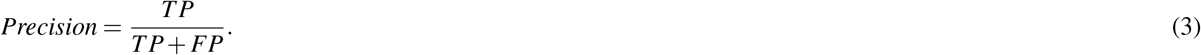

Numpy library^49^ was used to compute means and standard deviations. Percentages of cell divisions and cell death through time were computed as follows:

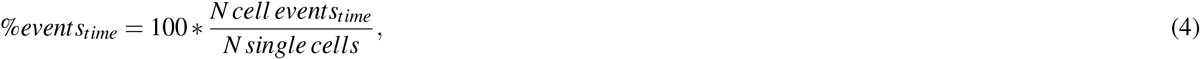

%*events*_*time*_ is the percentage of cell division or death at given time point; *N cell events*_*time*_ are number of cell divisions or death at given time; *N single cells* is the initial number of single cells at beginning of time-lapse. Division rate and death rate through time were computed upon a 6-hour temporal window:

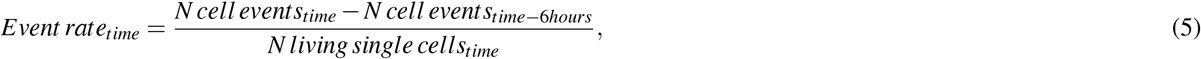

*Event rate*_*time*_ is the division or death rate at given time, *N cell events*_*time*_ are the number of cell divisions or death at given time, *N living single cells*_*time*_ is the remaining number of single cells that are still alive and have not divided yet at given time.

## Acknowledgements

GDR Imabio founded Hackathon Anger 2019 where AJC, SM and DR conceived study. AJC was funded by Hospices civils de Lyon and ITMO Cancer Soutien pour la formation à la recherche fondamentale et translationnelle en Cancérologie. The experiments where funded by a grant from the Ligue Nationale contre de le Cancer, comité Auvergne-Rhône-Alpes (MG and SM) and by a grant from Institut Convergence PLAsCAN, ANR-17-CONV-0002 (MG and SM). Charlotte RIVIÈRE kindly provided micro-fabricated chip and has to disclose the patent FR3079524A1.We thank Mrs Clarisse Hyron for english reviewing of the manuscript

## Author contributions statement

AJC, DR and SM conceived experiments. MG, OCE, DM and SM contributed to the study design. AJC conducted experiment, developed algorithms, annotated databases, is the main writer and made figures. AJC, DR and SM analysed data. AI and MG provided biological samples. AJC, NB, MG and CI cultured cells. All authors contributed to write and/or reviewed the manuscript and agreed to the published version.

## Additional information

AI reports research grants and travel funding from Carthera, research grants from Transgene, research grants from Sanofi, research grants from Air Liquide, travel funding from Leo Pharma, research grants from Nutritheragene, outside the submitted work.

